# BitterMatch: Recommendation systems for matching molecules with bitter taste receptors

**DOI:** 10.1101/2022.01.13.476205

**Authors:** Eitan Margulis, Yuli Slavutsky, Tatjana Lang, Maik Behrens, Yuval Benjamini, Masha Y. Niv

## Abstract

Bitterness is an aversive cue elicited by thousands of chemically diverse compounds. Bitter taste may prevent consumption of foods and jeopardize drug compliance. The G protein-coupled receptors for bitter taste, TAS2Rs, have species-dependent number of subtypes and varying expression levels in extraoral tissues. Molecular recognition by TAS2R subtypes is physiologically important, and presents a challenging case study for ligand-receptor matchmaking. Inspired by hybrid recommendation systems, we developed a new set of similarity features, and created the BitterMatch algorithm that predicts associations of ligands to receptors with ~80% precision at ~50% recall. Associations for several compounds were tested in-vitro, resulting in 80% precision and 42% recall. The encouraging performance was achieved by including receptor properties and integrating experimentally determined ligand-receptor associations with chemical ligand-to-ligand similarities. BitterMatch can predict off-targets for bitter drugs, identify novel ligands and guide flavor design. Inclusion of neighbor-informed similarities improves as experimental data mounts, and provides a generalizable framework for molecule-biotarget matching.

## Introduction

The sense of taste is a key driver in food choice and consumption^1^, and therefore has major implications for nutrition and health. The ability to taste is impacted in several diseases^2,3^, including COVID-19 infection^4^. Similarly to smell loss, taste loss or dysfunction can negatively affect the quality of life^5^ and promote changes in body weight^6^. Sweet, umami and bitter tastes are mediated by G protein-coupled receptors (GPCRs)^7^. Furthermore, taste GPCRs were shown to be expressed in many extraoral tissues, suggesting physiological roles beyond taste perception^8^, such as mediation of hormone secretion^9^, regulation of upper respiratory innate immunity^10^ and more.

Bitter taste receptors (TAS2Rs or T2Rs)^11^ present a particularly interesting case, where some TAS2Rs may be activated by tens of diverse ligands, whereas others are very selective and can be activated by only a few known ligands^12,13^. In addition, the number of TAS2R subtypes varies across species, with 25 in humans and ~30 in rodents^14^. Some bitter molecules activate several TAS2Rs while others are specific for individual TAS2Rs^13,15^. Bitter molecules have highly variable chemical structures, and include alkaloids, polyphenols, peptides, salts, fatty acids, and saponins^16^.

Addressing the dire challenge of predicting TAS2R targets for bitter molecules has several important implications. First, it can assist in evaluating off-targets for bitter medicines; since TAS2Rs have additional physiological roles in extraoral tissues, unintentional activation of ectopic TAS2Rs may promote unwanted biological processes as side effects^17^. Second, specific agonists for TAS2Rs provide tools for studying the extraoral effects of these receptors, or even as potential drug candidates targeting TAS2Rs for gastrointestinal^18^ and asthma indications^19^. Third, since numerous drugs and food compounds are intensely bitter, antagonists are needed to reduce bitterness and improve drug compliance, hence, designing antagonists for effective masking of bitterness relies on being able to identify the activated TAS2Rs^20–22^. Identification of TAS2R targets for a given molecule currently requires extensive in-vitro assays that consume time and resources, though some computational attempts to predict TAS2R targets began to appear^23^.

Computational methods are highly desired due to their high speed and low cost, as well as their potential of improvement as the experimental knowledge base grows. Specifically for bitter taste receptors, computational tools were developed for bitterness prediction using docking to homology models of selected receptors^21,24,25^ and ligand-based methods^16,26^. For the GPCR family at large, several machine-learning methods were developed to predict potential GPCR targets for small molecules, including machine learning algorithms^27,28^.

The problem of predicting associations of TAS2Rs and ligands can be viewed as a recommendation problem. Generally, recommendation systems are aimed at rating pairings of items to categories^29,30^. These correspond to ligands and receptors in our case. Content-based recommendation systems^31^ rely on attributes describing the items and the categories. However, collaborative recommendation systems rely on similarity measures from known associations or ratings^32^. Hybrid recommendation systems^33^ combine the two approaches by incorporating both content-based and collaborative information.

Here we develop BitterMatch, a classifier inspired by hybrid recommendation systems, to match bitter molecules to human and mice TAS2Rs. BitterMatch uses a novel set of features, which weight known associations between ligands and receptors by ligand-to-ligand and receptor-to-receptor similarities. Two BitterMatch scenarios are presented: *“filling the gaps”* for ligands for which some associations to TAS2Rs are known from cell-based experiments but some are missing, and *“new ligands”* for molecules for which bitter taste was established in human sensory tests, but no associations with individual TAS2Rs were measured.

## Results

### Ligand-receptor associations

Data on associations between bitter molecules and 21 human and 20 mouse TAS2Rs was collected from the literature to create the association matrix. The matrix is sparse, since some ligands were tested only on a subset of TAS2Rs. Out of 4501 known associations for 303 ligands (36% of possible associations) 3761 are negative (ligands not activating the receptor, ~84%) and 740 are positive (ligands activating the receptor, ~16%).

In agreement with previous observations^13^, TAS2R14 is the most broadly tuned human receptor, with 171 known agonists, followed by TAS2R39, TAS2R46 and TAS2R10 with 85, 79 and 48 ligands respectively. The most selective is TAS2R3 with only one known ligand. Mouse receptors (Tas2rs) also range from broadly tuned receptors, such as Tas2r105 (47 known agonists) to narrowly tuned Tas2rs, such as 122 and 139, each with one known ligand. In general, less screening experiments were performed for mouse Tas2s. Orphan TAS2Rs (that have no known ligands so far, four in humans and fourteen in mice) were excluded from training and inference. Tas2r113 was excluded due to having a single agonist that was not tested on human TAS2Rs.

### Ligand and receptors properties

For ligands, 2-dimensional (2D) and 3-dimensional (3D) chemical features including MW, AlogP, QPlogHERG were calculated from the chemical structures of the molecules.

For receptors, three types of chemical features were calculated: 1. Sequence-based features for the full protein sequence of the receptors and separately for the second extracellular loop, an important region for binding ligands in GPCRs^34^. 2. Binding site features that were calculated for the main site for ligands (the orthosteric binding site). 3. Structural features of the receptor, such as hydrophobicity and volume.

### Similarity calculations needed for features derivation

For ligands, we compute chemical similarity of the molecules using Tanimoto scores of the linear fingerprints, as well as Tanimoto scores of the MOLPRINT2D fingerprints.

For receptors, we compute sequence similarities based on the percent of identical positions in the protein sequences (percentage of sequence identity), and in the sub-sequences that consist of the orthosteric binding site residues. Additionally, since substitution between non-identical residues are well studied^35^, we calculate similarities between the sequences of the receptors and the sub-sequences of the orthosteric binding site based on BLOSUM62 substitution matrix^36^ (percentage of sequence similarity). All similarities are schematically represented in Figure 1A.

**Figure 1.**
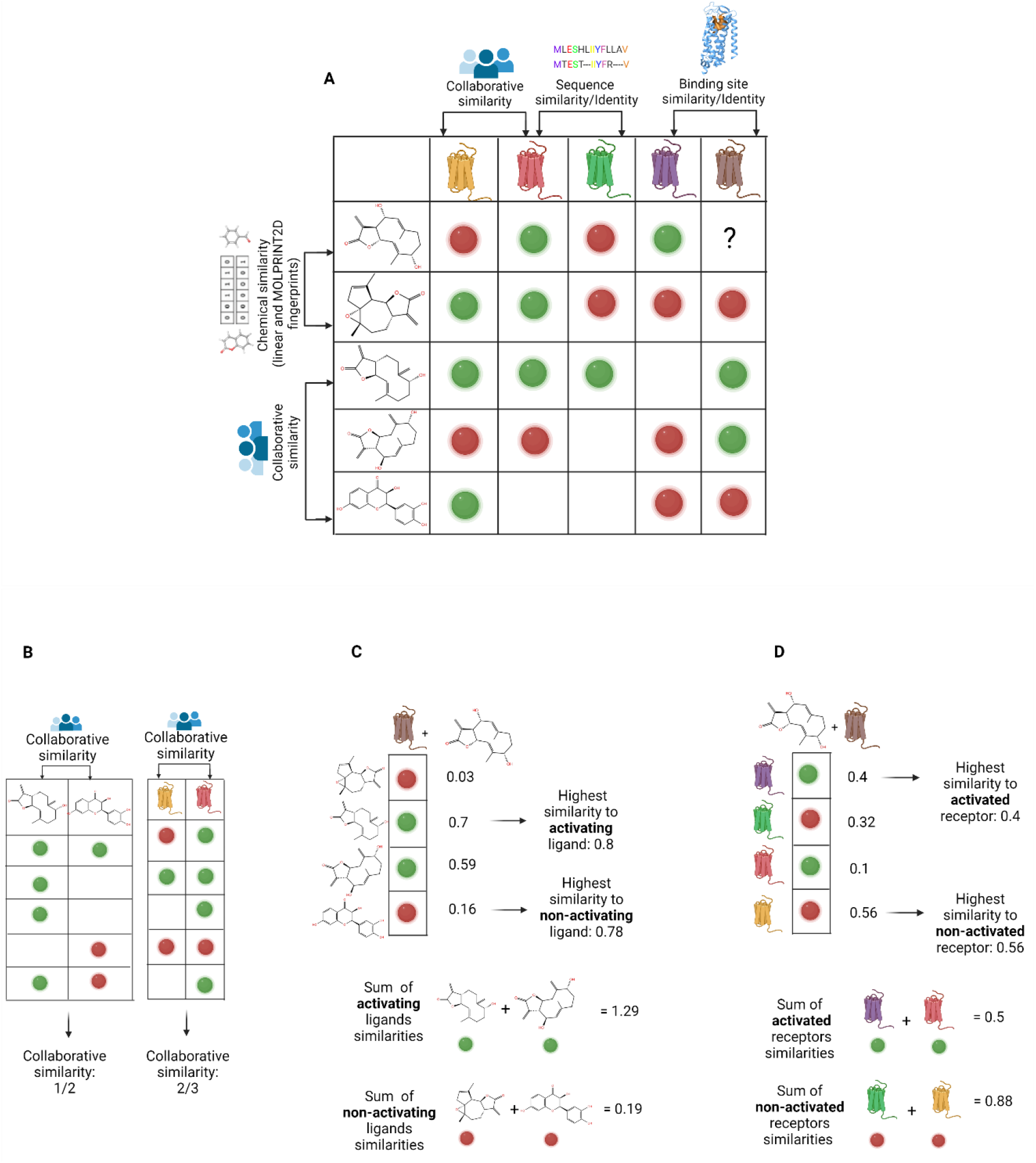
Similarity types and similarity-based features. **A** –Ligand similarities include chemical similarities based on linear fingerprints, chemical similarities based on MOLPRINT2D fingerprints, and collaborative similarities. Receptor similarities include collaborative similarities, similarities calculated from sequence identity matrices, and from sequence similarities derived from BLOSUM62 substitution matrix. Sequence similarities were calculated based on the full protein sequence and based on the binding site sequences. Colored circle: red – the ligand does not activate the receptor, green – the ligand activates the receptor. Unknown associations are represented as blank spaces in the matrix. **B** – Collaborative similarities for pairs of ligands and pairs of receptors, calculated based on known associations using Jaccard similarity. **C** – Four similarity features are computed for ligands: the highest similarity to the ligand that activates *r*, sum of similarity values to all ligands that activate *r*, the highest similarity to the ligand that does not activate *r*, and the sum of similarity values to all ligands that do not activate *r.* **D** – Similarity based features are computed for the receptor: the highest similarity to the receptor that is activated by *l*, the sum of similarities to receptors that are activated by *l*, the highest similarity to the receptor that is not activated by *l*, and the sum of similarities to receptors that are not activated by *l*.

Collaborative similarities between pairs of ligands (or receptors) were calculated as Jaccard similarities over their known associations, and used to construct ligand-to-ligand and receptor-to-receptor collaborative similarity matrices (Figure 1B).

### Extracting neighbor-informed features from similarities

To avoid dependence of the learning algorithm on the size of the dataset, we devised new neighbor-informed features, which are based on the similarity matrices and the known associations. The features are coded separately for positive and negative associations. Specifically, from a ligand similarity matrix, we annotate each ligand (*l*)-receptor (*r*) pair with four features: two summarize the similarities of *l* to ligands with positive associations to the receptor *r* and two summarize the similarities of *l* to ligands with negative associations to *r*. The features differ by their granularity: the first measures the similarity to the closest ligand that activates *r*; this feature represents positive examples in the local neighborhood. The second measures the sum of similarity values to all ligands that activate *r*. The two negative features measure the similarity to the nearest ligand that does not activate *r*, and the sum of similarities of ligands that do not activate *r* (Figure 1C). We repeat this feature extraction process for each ligand similarity matrix.

We also extract neighbor-informed features from each receptor similarity matrix, reversing the roles of ligand and receptor: similarity to the closest receptor that is activated by *l*, the sum of similarities to receptors that are activated by *l*, similarity to the closest receptor that is not activated by *l*, and the sum of similarities to receptors that are not activated by *l* (Figure 1D).

### *“Filling the gaps”* scenario

The association matrix between bitter ligands (n = 303) and human or mouse TAS2Rs (m=21+20) has about 1/3 known and 2/3 unknown associations. In *“filling the gaps”* scenario we consider cases in which at least one association (positive or negative) is known for each ligand and for each receptor, and therefore, neighbor-informed features from the similarity matrices can be extracted for ligands and receptors.

We model the problem as a binary classification task, in which each ligand (*l*)-receptor (*r*) pair is considered an observation and is annotated with (a) features describing chemical properties of *l* (*p_l_* = 250), (b) features describing the chemical properties of *r* (*p_r_* = 235), (c) features derived from the similarities between *l* and other ligands (*p_sl_* = 4 features o 3 types of similarities) and features derived from the similarities between *r* and other receptors (*P_sl_* = 4 features · 5 types of similarities). We train a classifier from these features using a gradient boosting algorithm with decision-tree learners (XGBoost^37^ package) optimizing a binary logistic objective for predicting whether the ligand-receptor pair associates.

### Evaluation of the model

We sample 80% of the known positive and negative associations to be used as a training set, and use the remaining 20% as a test set. We repeat this sampling process 100 times. The performance of BitterMatch is compared with a naive model that predicts for each ligand-receptor pair whether they associate according to the prior. In the prior, the prediction score is fixed per receptor for all ligands and is set to the proportion of ligands in the training set known to associate with the receptor. We further compare the full model with three submodels that contain different subsets of the features, as illustrated in Supplementary Figure S1. Model 1 includes only chemical properties of ligands and receptors; Model 2 includes in addition also features based on collaborative similarities; Model 3 includes chemical properties as well as neighbor-informed chemical similarity features. Model 4 is the full model.

Average precision-recall curves over the repetitions are reported in Figure 2 (Supplementary Figure S2 shows prediction intervals). Model 1, which uses only chemical properties of ligands and receptors, already performs better (average precision 65% ± 2 %) than the prior (49% ± 0.5%). Adding collaborative similarity-based features (Model 2) further improves the performance (71% ± 2%). Adding features that incorporate both known associations and chemical and sequence similarities leads to an even larger improvement (Model 3, 76% ± 3%). Model 4, in which collaborative features are added as well, performs as well as Model 3, which is therefore the chosen model. For Model 3, a threshold *t* = 0.5 on the predicted probabilities for each ligand-receptor pair yields precision of 84% and recall of 51% on average over the 100 repetitions.

**Figure 2.**
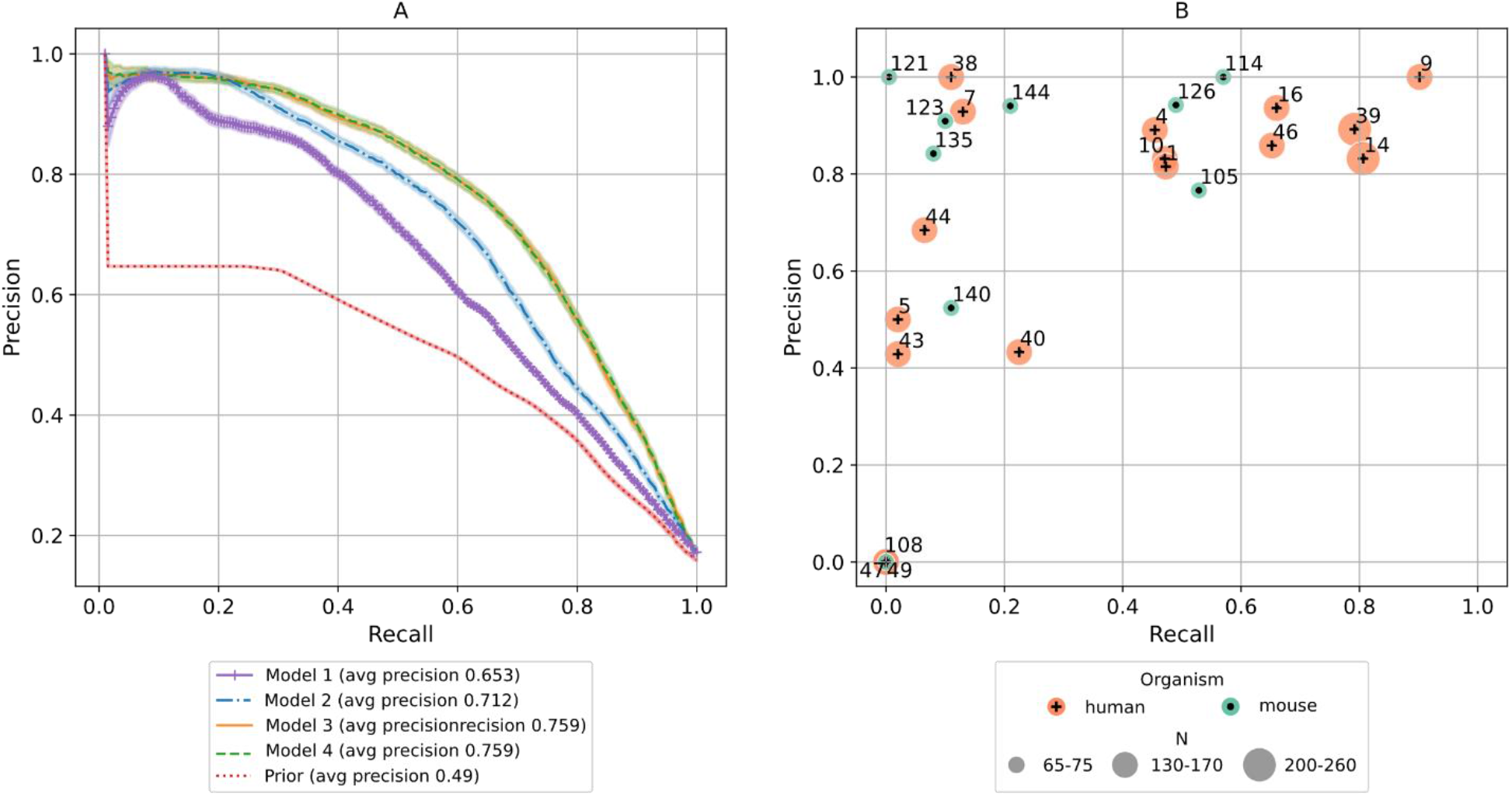
BitterMatch performance for filling the gaps. **A** – Average recall-precision curves over 100 repetitions of the experiment for the prior model and 4 BitterMatch sub-models. 95% Bootstrap confidence intervals for the mean are shown for each model. **B** – Average recall and precision per receptor according to predictions of the recommended Model 3. Each point represents a receptor; human receptors are shown in orange, mouse receptors in green. The point sizes represent the number of associations known for the receptors. Receptors for which no positive predictions were made are omitted from the figure.

Recall and precision levels per receptor averaged over 100 repetitions, are shown in Figure 2B. The performance is generally better for receptors with more known (positive and negative) associations. Performance is higher for human receptors than for mice receptors, in accordance with a higher number of known positive associations. Nevertheless, mouse receptors 112, 126 and 105 all achieve recall above 49% and precision above 76%.

### *“Filling the gaps”* predictions

Using BitterMatch Model 3 we filled the gaps in the sparse association matrix, and predicted the unknown association of the 7,922 missing pairs of bitter compounds and human and mouse TAS2Rs (Figure 3). We used a threshold of 0.65 (prioritizing high precision). The experimental data that was used for training and testing was supplemented by the new predictions to construct a full association matrix without gaps. Analysis of the complete association matrix at the chosen threshold revealed that out of 12,423 associations, 1479 are positive (12%) and 10,944 are negative (88%). The predictions are available via the [BitterDB](http://bitterdb.agri.huji.ac.il/dbbitter.php) database. As expected, TAS2R14 is still the most broadly tuned receptor with 191 ligands (171 known and 20 predicted ligands). Some receptors, such as TAS2R40 and Tas2r121, had no new positive predictions, while others gained many newly-predicted positive associations (Figure 3).

**Figure 3.**
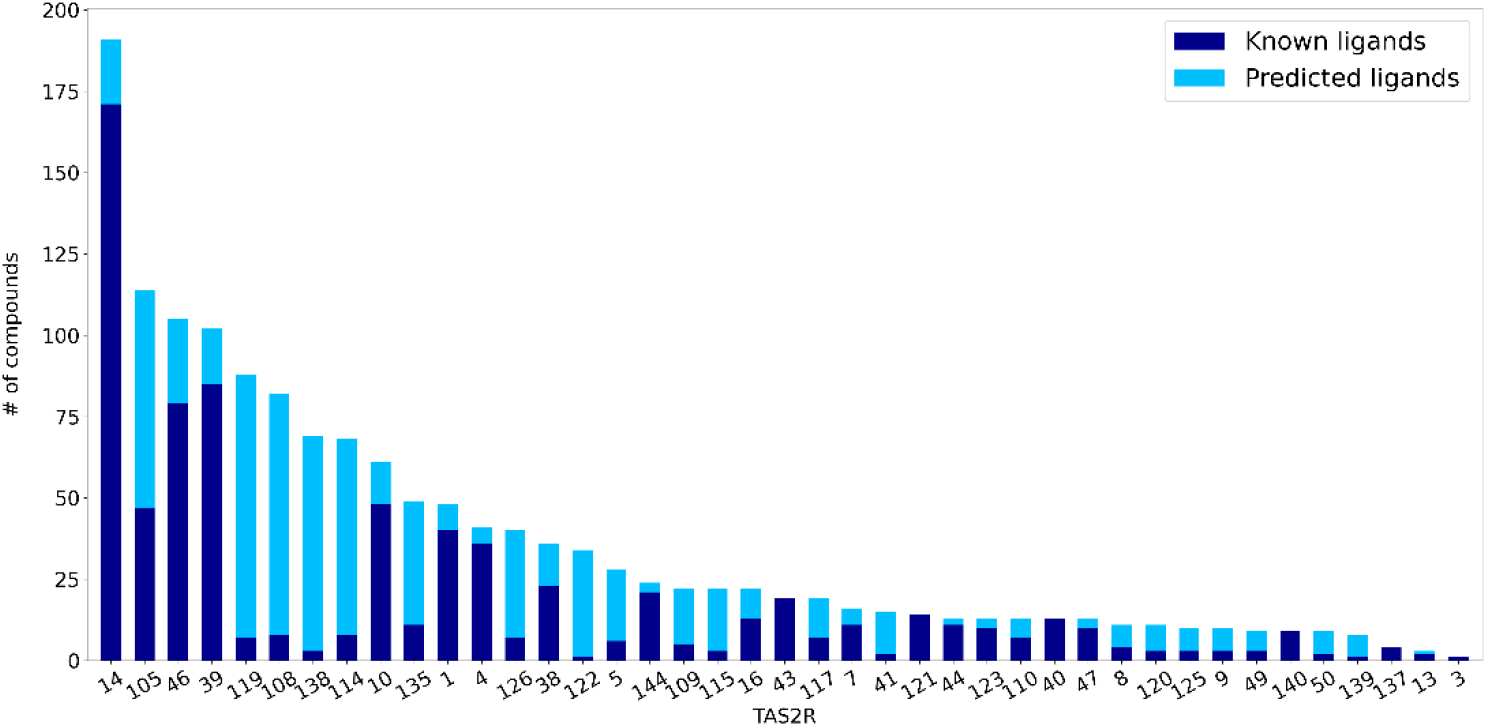
Filling the gaps with new predicted associations. Names of TAS2Rs (digits for human, hundreds for murine) ordered by decreasing number of ligands. The number of experimentally validated ligands per TAS2R is presented in dark blue (known ligands). The number of predicted ligands is presented in light blue.

Since different species have varying numbers of TAS2R subtypes^38^, assigning TAS2R functional analogs is not trivial. This task is especially important for humans and mice, since many taste examinations are performed on rodents, but aiming to reflect on humans^39^. In order to find the nearest functional receptor (“functional analogs”), we used the completed association matrix to calculate the Jaccard similarity between the receptors, based on their positive and negative, known and predicted associations. No score above 0.5 between human and mouse TAS2Rs was found (see supplementary data), suggesting relatively low overlap and no clear functional analogs.

### *“New ligands’’* scenario

In this scenario we predict associations with human TAS2Rs for bitter molecules that do not have any known association with bitter taste receptors. Collaborative similarities, as well as features based on neighbors-informed receptor similarities, can not be calculated in this case. Therefore, we develop a version of BitterMatch that uses only the chemical properties of ligands and of receptors, and neighbors-informed **ligand** similarity features.

For evaluation of the performance, we sample 80% of the ligands in the dataset into a training set, and consider the remaining 20% as a test set. For the ligands in the test set we remove the associations with all the receptors, marking them as unknown, and repeat this process 100 times. We compare our model to a prior model (as in the previous section), and to a “nearest-neighbor” model that predicts association between a ligand *l* and a receptor *r* based on the known association of *r* with the ligand that has the highest chemical similarity with *l*. Here chemical similarity is calculated according to linear fingerprints (using MOLPRINT2D fingerprints yielded similar results, not shown).

The average precision-recall curves over 100 repetitions and corresponding confidence intervals are reported in Figure 4A (prediction intervals are shown in Supplementary Figure S3). BitterMatch achieves an average precision of 70% ± 5%, outperforming the prior model (average precision of 48% ± 4%) and the nearest-neighbor model (44% ± 5%).

**Figure 4.**
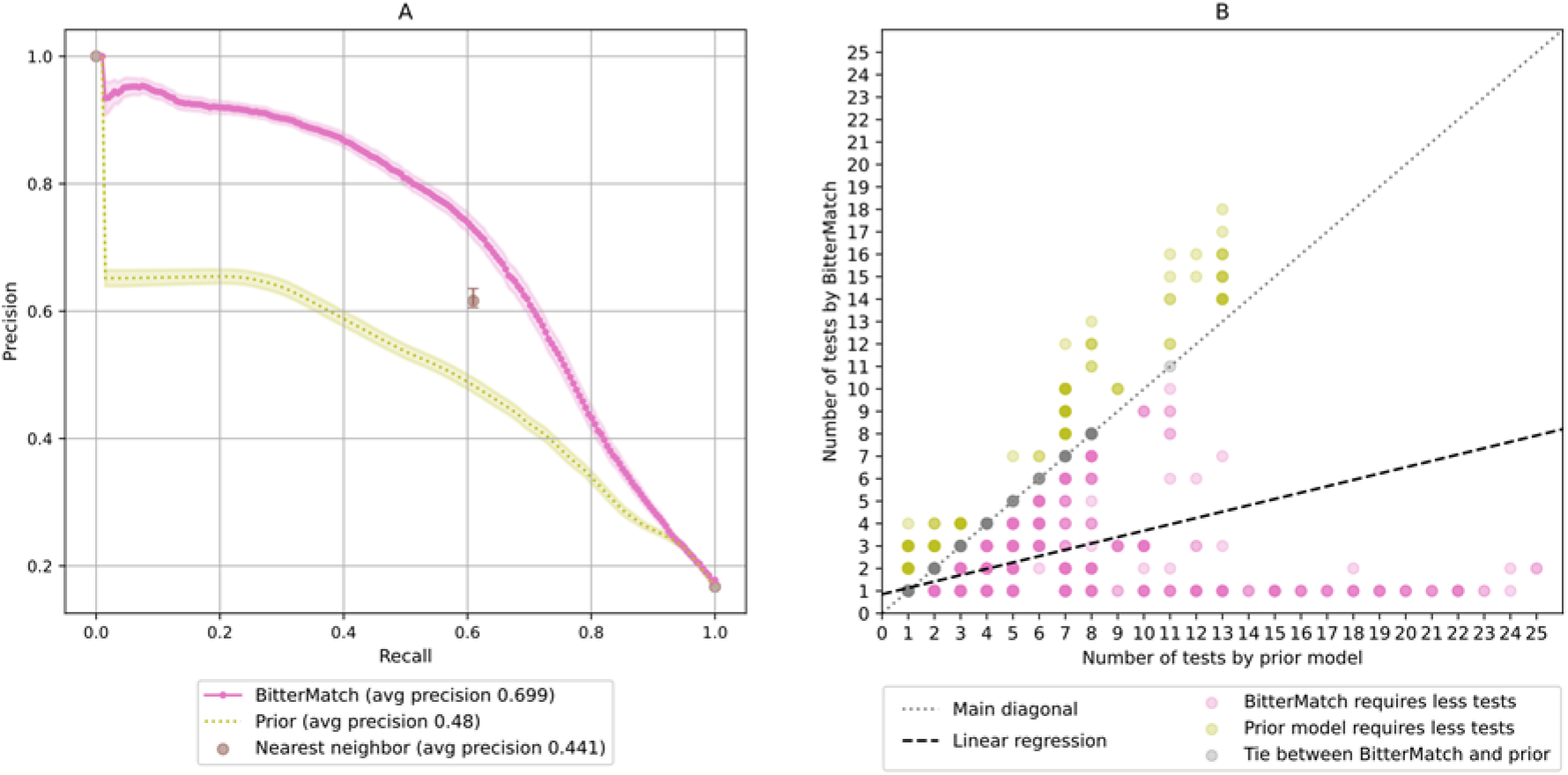
Performance of “new ligands” BitterMatch. **A** – Average recall precision curves over 100 repetitions for the BitterMatch model, a prior model and a nearest neighbor model. 95% Bootstrap confidence intervals for the mean are shown for each model. **B** – The number of tests required to find at least one receptor which is activated by the ligand. Each point represents a ligand in one of the test sets in 100 repetitions of the experiment. Experiments in which BitterMatch required less tests are shown in pink below the main diagonal, experiments in which the prior model required less tests are shown in yellow above the diagonal. Experiments in which the two methods required the same number of tests are shown in grey on the main diagonal. Darker points correspond to more experiments. The black dashed line is a fitted linear regression (for the number of tests by BitterMatch as a function of the number of tests by the prior) with intercept of 0.852 and coefficient of 0.283.

An accurate predictor of positive association can reduce the number of cell-based experiments needed to identify cognate bitter taste receptors for a bitter molecule. Therefore, for each ligand we count how many receptors should be tested, according to the prediction score, until the first TAS2R is activated. Figure 4B shows the number of tests required by the prior method and by BitterMatch. To quantify the differences, we fitted a linear regression that predicts the number of tests required by BitterMatch as a function of the tests required by the prior. On average, BitterMatch requires 3.5 times less cell-based experiments in order to match at least one TAS2R to a bitter-tasting molecule without any known bitter taste receptors.

### Prospective predictions and their validation

To evaluate the model in a prospective mode, we predicted the associations of 12 new bitter compounds. The validation set included 3 new compounds and 5 compounds from BitterDB^12^ for which only a few associations were known and were excluded from the original association matrix. Functional screening of the 21 human TAS2Rs expressed in HEK 293T-Gα16gust44 cells with the 8 proposed agonistic compounds resulted in the identification of receptors for all of the substances. After the initial identification of presumed receptor-compound pairs, all compounds were tested with three concentrations on the identified receptors to assess their potencies (Fig. S5 in the Supplementary data). The lowest concentrations leading to statistically significant activation of receptor-expressing cells compared to mock-transfected cells were judged as apparent threshold concentration. All 8 compounds resulted in the activation of TAS2R14. Whereas 2-acetyl benzofuran, butein, 3,2’-dihydroxychalcone and sinapic acid exclusively activated the TAS2R14, the other four compounds elicited responses of at least one additional TAS2R: Apigenin activated the TAS2R43, theacrine the TAS2R43 and TAS2R46, and fisetin and quercetin activate TAS2R4, -R7, -R10, -R31, -R39, -R40, -R43, and -R46. The highest signals were observed for TAS2R14 when activated by quercetin and the lowest activating concentrations were 0.3 μM (butein, TAS2R14; fisetin, TAS2R7, -R10, -R14, -R40, -R43; quercetin, TAS2R14). The results are summarized in Suppl. Table S1. We note that for butein, 3,2’-Dihydroxychalcone and apigenin activation of TAS2R39 was not detected in our experiments but was found by Roland *et al*.^40^, hence we present the results with and without these three associations. BitterMatch predictions for 4 additional compounds were compared to the recently published cell-based measurements.^41^

For the 12 compounds in total, the model correctly predicted 16 true positive pairs and 210 true negative pairs, but misclassified 4 pairs as positives and 22 pairs as negatives (Table 1), hence we achieved precision of 80% and recall of 42% on the validation set. By excluding three inconsistent associations with TAS2R39 (as described above), we achieve precision of 76% and 37% recall.

**Table 1.**
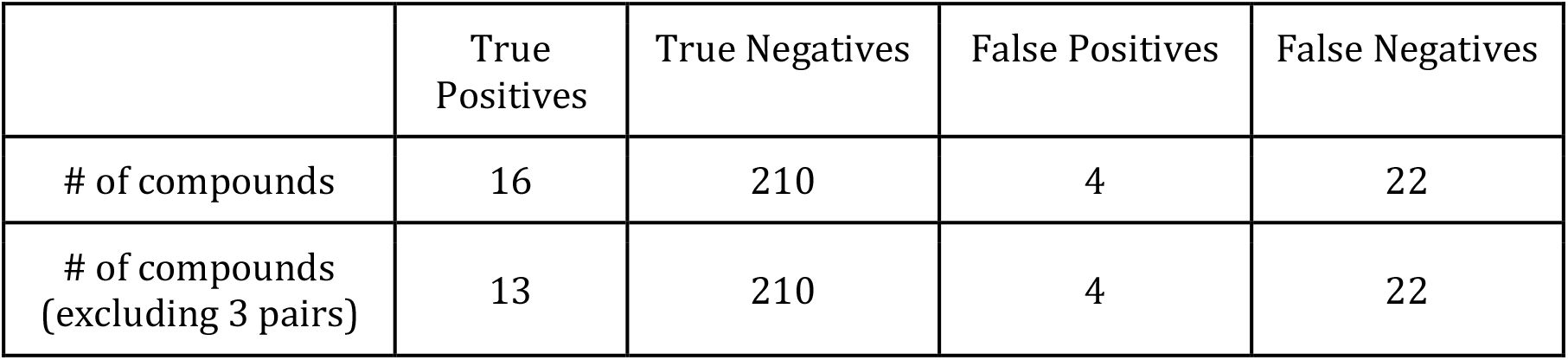
BitterMatch results on the validation set.

Three out of four false positive pairs were wrongly assigned to TAS2R14, the most broadly tuned receptor. False negatives stemmed from 8 missed associations with TAS2R43, 4 with TAS2R46 and 2 with each of TAS2Rs: 4,7,10,40 and 44, details in Supplementary Table S3.

### Predicting TAS2R associations of drugs from DrugBank

As a possible application, we used BitterMatch to predict associations of drugs (DrugBank dataset version 5.1.5)^42^ with TAS2Rs. 2406 out of 10,170 compounds were predicted to be intensely bitter using the BitterIntense algorithm^16^.

For those, BitterMatch predicted 2,207 positive pairs to TAS2Rs: 1576 for TAS2R14, 394 for TAS2R46, 127 for TAS2R10 and the others for TAS2Rs: 1,4,7,16,38 and 39 (Figure 5). Comparing the proportion of drugs matched to TAS2R with the proportion of positives among all tested compounds for that TAS2R (hit rate) suggests that the proportion of drugs activating similar to hit rate for TAS2R14 but much lower for TAS2Rs 10, 39 and 46. The results strengthen the notion of importance of TAS2R14 as a major target of bitter pharmaceuticals^43^ (Figure 5).

**Figure 5.**
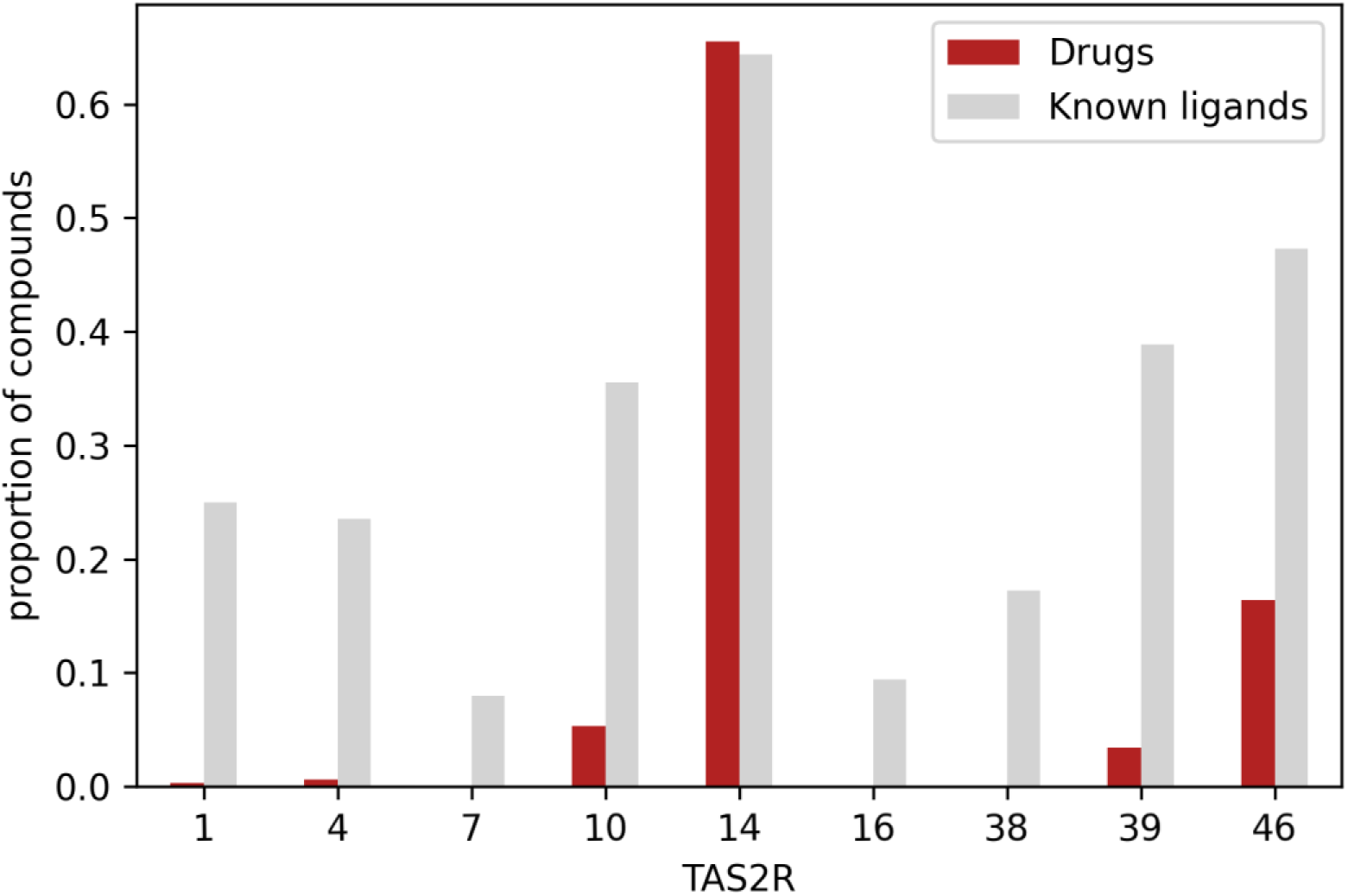
Matching intensely bitter drugs to human TAS2Rs. Very bitter predicted drugs from DrugBank (version 5.1.5) were assigned to human TAS2Rs using BitterMatch. The proportion of drugs that were predicted to activate each receptor represented in red. In grey-the proportion (hit rate) of known ligands per receptor. Normalization for the drugs was performed by dividing the number of drugs predicted to activate the receptor by the total number of drugs. The hit rate was calculated by dividing the number of ligands by the number of compounds that were tested for each specific receptor.

### Feature importance

To obtain intuitive insights into the model, we examine the average gain across all XGBoost splits (Methods: “Feature importance”). This analysis shows few features showing considerably higher importance values than others (Figure 6 and Supplementary Figure S4). In both *“filling the gaps”* and *“new ligands”* scenarios, neighbor-informed features are most important: similarities of the ligand to both activating and non-activating ligands of the receptor. Receptor neighbor-informed features could not be calculated in “new ligands” scenario, but found to be important for *“filling the gaps”.* In both scenarios, receptor chemical properties are important, top ones include hydrophobicity, charge and buried area properties. Ligand properties have low gain in both scenarios (Supplementary Figure S4).

**Figure 6.**
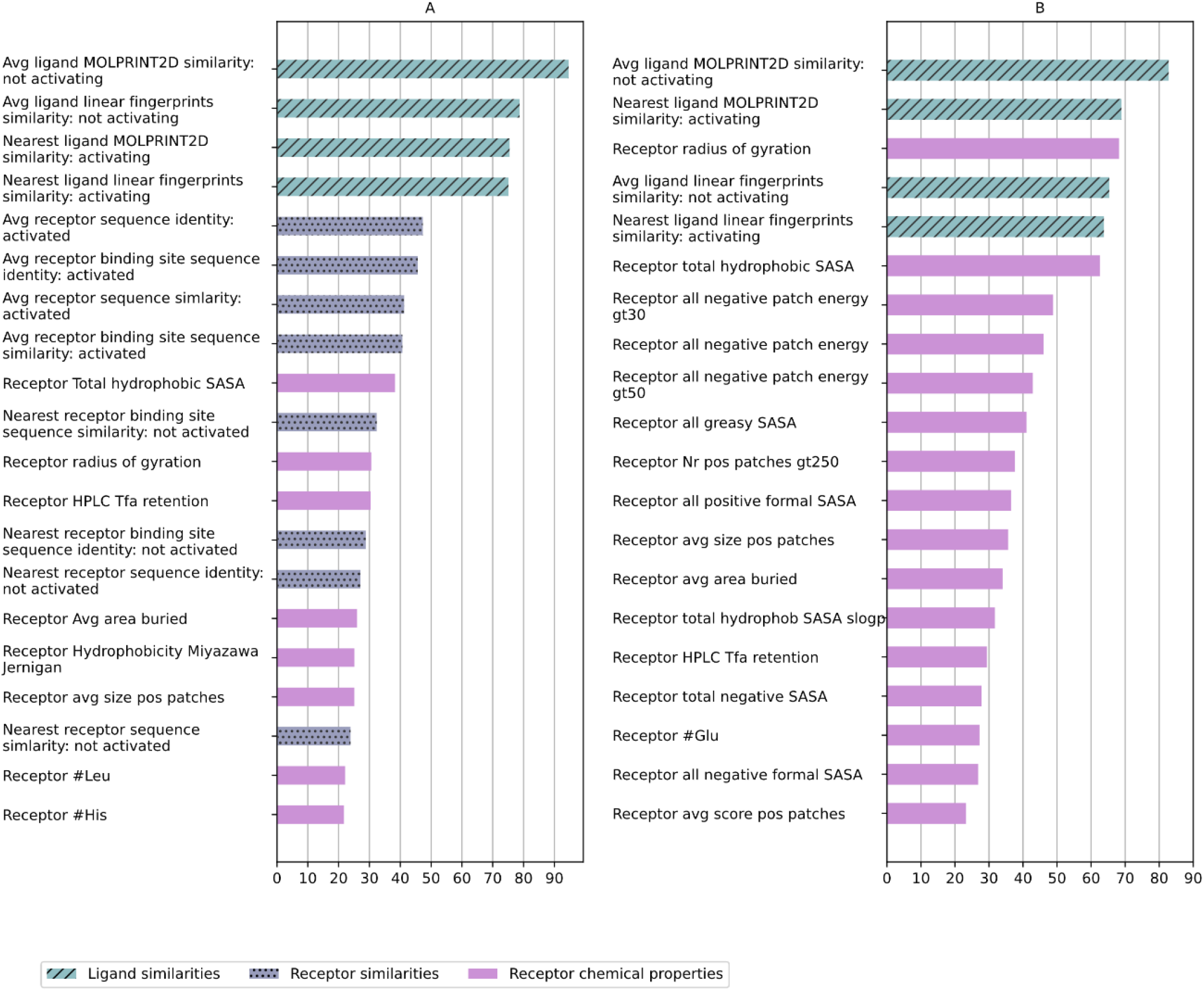
Feature importance for filling gaps. Features related to ligand similarities are shown in blue with diagonal lines, features related to receptor similarities (included only in *“filling the gaps”* model) are shown in purple dotted bars. Receptor properties are shown in plain purple. (A) – Feature importance of top 20 features of model 3 (corresponding to average gain over 21.5). The y axis corresponds to features (ordered by importance) and the x axis shows the average gain over XGBoost splits. (B) – Feature importance of top 20 features of the *“new ligands”* model (corresponding to average gain over 22.7)

## Discussion

Matching ligands with their receptors is at large an open task in biorecognition and in drug discovery. For TAS2Rs, it is especially challenging due to the dramatic chemical diversity of bitter ligands, the high variance in the number of receptors that each ligand activates, and the lack of experimental structures of the receptors.

In this work we presented BitterMatch, an algorithm designed to predict ligand-TAS2R associations. BitterMatch was modeled as a binary classification task, with a custom feature set composed of chemical descriptors as well as neighbor informed features extracted from multiple similarity matrices. We chose a tree boosting algorithm (XGBoost) as the learning algorithm due to its relative success with unbalanced classes and sparse.

Despite the challenges, we achieved promising results with average precision of 70-76% for both *“filling the gaps”* and *“new ligands”* scenarios. Importantly, the high precision and recall were not only established in the train-test divisions of the dataset, but also confirmed in prospective predictions. Three associations predicted as positives for TAS2R39, were found positive in^40^ but negative in our experiments. This could be due to stable vs. transient expression of the TAS2R gene, which might affect the sensitivity of the cells toward activation. Excluding these 3 associations from the analysis resulted in precision of 76% and recall of 37% instead of 80% and 42%.

The novel neighbor-informed similarity-based features, which incorporate chemical similarities and information from known positive and negative associations, dramatically improve the performance of the algorithm. In contrast, chemical features of the ligands do not provide high gain, in accordance with the high chemical diversity of bitter molecules, which makes it difficult to associate specific chemical attributes to the ability of a molecule to activate specific TAS2Rs. On the other hand, chemical features of the receptors, in particular those related to hydrophobicity and net charge, substantially contribute to the model. This is in accord with hydrophobicity of the binding site importance for enabling recognition of multiple ligands^13^. The fact that orthosteric binding-site similarities play a dominant role in *“fill the gap”* scenario, suggests that this is the site most of the ligands bind to^44,45^. The importance of the overall similarities of receptors suggests that access to the binding site is likely to play a key role as well^45^.

In addition, we show that combining the chemical features of both ligands and receptors together with neighbor-informed ligand similarity features leads to much higher performance than relying only on the known associations of the most similar ligand, or on the number of receptor’s positive associations.

Rodents are used as a model for bitterness assessment, sharing a similar repertoire of bitter compounds^46,47^. Our results suggest that at the individual receptor subtype level, activation of a rodent receptor is not predictive of the human receptor and vice versa. This is particularly important when using animal models to elucidate the physiological roles of extra-oral TAS2Rs^48^.

Computational matching of TAS2Rs and their agonists by BitterMatch can assist in identifying specific agonists for TAS2Rs, elucidating functional relations between TAS2Rs and evaluating the potential TAS2R targets of food and drug compounds. We used the model to predict associations of intensely bitter drugs to human TAS2Rs. The results revealed that TAS2R14 is the main receptor in pharmaceutical drugs bitterness. TAS2R14 is known to be activated by antibiotics^49^ and other diverse drugs^43^. It was also suggested that TAS2R14 regulates resveratrol transport across the human blood-cerebrospinal fluid barrier^50^.

While BitterMatch yields good performance overall, limitations of the method should be kept in mind. The performance of BitterMatch is worse for receptors for which less associations are known, and in particular those with less positive ones. Indeed, since the similarity-based features also depend on known associations, in *“filling the gaps”* task we achieved poor results for ligands with few experimentally known positive associations. We expect further improvement as more associations are gathered experimentally. Additionally, since each TAS2R was represented by a single protein sequence (according to the common allele), mutations or variations in the sequence that might alter ligand recognition or receptor activation were not taken into account.

Unraveling the associations of bitter compounds with TAS2Rs will advance studies of extraoral TAS2Rs, shed light on involvement of TAS2Rs in health and disease and in finding chemosensory relations across species. Incorporation of BitterMatch-like methods in drug discovery may enable computational prediction of off-targets, and save in-vitro experiments, time and resources.

The novel neighbor-informed similarity features can be generalized to other GPCRs and other small molecule-biotarget matching.

## Methods

### Data curation

We curated a dataset consisting of known associations for pairs of TAS2Rs and bitter molecules from BitterDB^12^ and other publications^21,40,51–53^. We considered a ligand-receptor pair to be positive (positive association) if the molecule activates the receptor. If the molecule does not activate the receptor, we considered this pair as negative (negative association). Only associations that were confirmed in *in-vitro* assays were considered. Associations that were not confirmed *in-vitro* or were inconclusive between different papers were considered unknown. The constructed association matrix included only agonists as true positives and did not include inverse agonists or antagonists. In addition, we did not use human and mouse receptors that are considered orphans (have no known ligand). Mouse Tas2r113 was excluded due to the fact that it has only one agonist that was not tested on human TAS2Rs. In total, for 21 human and 20 mouse TAS2Rs and 303 ligands, we collected 4501 pairs of ligand-receptors out of which 740 are positive associations and the 3,761 negative associations.

## Ligand features

### Ligand preparation

After obtaining SMILES strings of the ligands from BitterDB and associations from the papers mentioned in “Data curation”, the compounds were uploaded to Maestro (Schrödinger Release 2019-2: MS Jaguar, Schrödinger, LLC, New York, NY, 2019). 3D structures of the ligands were generated using Ligprep and Epik (Schrödinger Release 2019–2: LigPrep, Epik, LLC, New York, NY, 2019) in pH 7.0 ± 0.5. In case the structures had additional fragments, these were removed. All compounds were desalted when possible, retaining the bigger ion and removing the smaller counter ion. Original chirality of compounds was retained when specified, otherwise additional stereoisomers per ligand were generated. For each ligand, the conformer with the lowest energy was chosen. When two stereoisomers were generated for a given compound, both structures were kept.

### Ligand chemical properties calculation

Three sets of descriptors were calculated for the prepared 3D structures using Canvas (Schrödinger Release 2019–2: Canvas, Schrödinger, LLC, New York, NY, 2019): Physicochemical descriptors, Ligfilter descriptors (moieties, atoms and functional groups) and QikProp descriptors (ADME descriptors). In total, 235 features were calculated. For the QikProp descriptors, additional PM3 properties were calculated as well (Schrödinger Release 2019–2: QikProp, Schrödinger, LLC, New York, NY, 2019). Several compounds that could not be neutralized in the ligand preparation process didn’t contain QikProp descriptors due to the limitations of calculating QikProp descriptors, however they were also included in the data set due to low amounts of true positives in the data. The full list of features can be found in the supplementary data.

## Receptor features

### TAS2Rs structure preparation

21 non orphan human TAS2Rs and 20 non-orphan mouse receptors were used. Pre-computed homology models were extracted for 21 human TAS2Rs and 14 mouse TAS2Rs from BitterDB. Additional 7 models were created using the I-TASSER server^54^. All the models were prepared and energy-minimized using the protein preparation wizard in Maestro (Schrödinger Release 2019–2) at pH=7±0.5 and default configuration.

### Receptor descriptors calculation

We calculated 3 sets of descriptors for the bitter taste receptors: 1. Sequence based features. 2. Binding site features 3. Structural features. The full list of features can be found in the supplementary data.

### Sequence based features

The set of sequence-based features was calculated (a) for the whole receptor and (b) for the extracellular loop number 2 (ECL2) separately, resulting in a total of 66 features per receptor. The protein sequences of the receptors were taken from BitterDB^12^ and uploaded to ProtParam^55^. The ECL2 sequences were extracted using TOPCONS server^56^ and uploaded to ProtParam. The ProtParam server calculated 33 features for each receptor, including the counts of the amino acids, molecular weight, theoretical isoelectric point (pI), count of negatively and positively charged residues, count of atoms, Grand average of hydropathicity (GRAVY), and Instability index. The full set of features are described in the documentation of ProtParam.

### Binding site features

The binding sites of the receptors were visualized and evaluated using the SiteMap tool in Maestro (Schrödinger Release 2019–2). Briefly, potential binding sites were shown in Maestro as a set of site points at or near the surface of the receptor that are contiguous or are separated in solvent-exposed region by short gaps that could plausibly be spanned by ligand functionality^57^. For each receptor, SiteMap suggested several pockets that could serve as binding pockets. Of the potential binding pockets sites that were oriented at the extracellular surface of the receptor, we chose the site with the highest scores, which usually correlates with the real binding sites^58^. We extracted 16 features from the binding sites: total volume, hydrogen bond acceptor volume, hydrogen bond donor volume, hydrophilic volume, hydrophobic volume, surface volume, binding site score, size, Druggability score (Dscore), hydrophobic and hydrophilic properties, exposure, enclosure, hydrophobicity, contact, balance, donor/acceptor.

### Protein 3D features

We calculated 166 protein descriptors including volume, exposed aggregation surface area, formal charges, area buried, size of positive and negative patches and more. Calculations used the “calc_protein_descriptors.py” script from BioLuminate package (version 3.5, Schrödinger, LLC, New York, NY, 2019) in Maestro.

### Additional information on the receptor

we added the chromosome number and the organism (human/mouse) to the feature set. This data was also taken from BitterDB^12^.

## Chemical similarities of ligands and receptors

We compute multiple similarity matrices for ligands and receptors. Each ligand is compared to another ligand (and receptor to receptor) creating *LxL* and *RxR* matrices.

### Ligand similarities

Two similarity matrices were generated for the bitter molecules in our data using two types of fingerprints. The first encodes a combination of defined linear fragments and ring closures; the second is extracted from the MOLPRINT2D fingerprint, a radial-like fingerprint that encodes atom environments using lists of atom types located at different topological distances, using Canvas (Schrödinger Release 2019–2). The calculation of fingerprints is encoded by hashing a defined set of sub-structures or chemical patterns into an N-bit address space, where only “*on*” bits are stored (meaning the particular sub-structure or atom appears in the molecules). The sub-structures or chemical patterns change according to the fingerprint type. After computing the fingerprints, the similarity between two molecules can be determined by comparing the *on* bits of the two structures. Denoting *on* bits in structure 1 by a, *on* bits in structure 2 by b and *on* bits in both structures by c, we take the Tanimoto similarity as defined by:

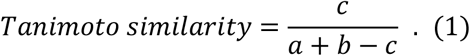

### Receptor sequence similarities and identities

Multiple Sequence Alignment (MSA) of human TAS2Rs was taken from BitterDB^12^. Mouse receptors were added into the alignment using ClustalW^59^ in Jalview^60^. The MSA was also validated by checking the alignment of conserved residues according to Di Pizio et al.^61^ Then, sequence identity (percentage of identical amino acids in aligned positions) and sequence similarity scores (calculated using BLOSUM62 substitution matrix^36^) were calculated for each pair of receptors using SIAS webserver (http://imed.med.ucm.es/Tools/sias.html) in default configuration. The results were in agreement with Chandrashekar et al.^11^ and Adler et al.^7^

### Receptor binding site: sub-sequence similarities and identities

To compare between the residues of the binding sites, all the residues within 4Å or 5Å from the docked flufenamic acid in the homology model of TAS2R14 were considered. The residues in the same positions for all the other receptors were extracted from the MSA. The matrices for subsequence similarity and identity were then calculated using SIAS as explained in “Receptor sequence similarities and identities”, for this subset of positions. Cutoff distance of 4Å led to decreased performance, thus 5Å distance cutoff was kept. TAS2R14 and flufenamic acid were chosen since flufenamic acid is a strong agonist of TAS2R14 and the interactions are well known from mutagenesis studies^21^.

## Collaborative similarities

In addition to pre-computed similarities, we construct two collaborative similarity matrices, one for the ligands (*S*^Lig^ ∈ ℝ^*L×L*^) and one for the receptors (*S*^Rec^ ∈ ℝ^*R×R*^), based on the associations. Denote by A∈ {0,1,na}^*L×R*^ the matrix of associations between *L* ligands (rows) and *R* receptors (columns). An entry *A_lr_* equals to 1 if the ligand *l* activates the receptor *r*, 0 if it is known not to activate the receptor *r*, and “na” if the association is unknown or removed for training purposes. We define the collaborative similarity in each of these two matrices as proportion of matching associations. For ligands *I* and *l*’ we define the collaborative similarity as

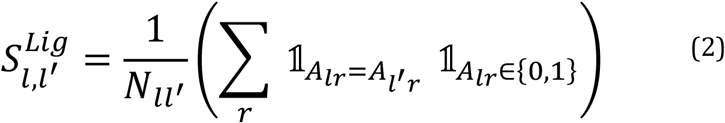

where *N_ll_*, is the number of receptors that their association with both ligands *l, l*’ is known.

Similarly, the collaborative similarity between two receptors, *r* and *r*’ is defined as

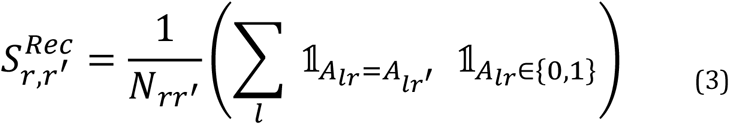

where *N*_*rr*_, is the number of ligands that their association with both receptors *r, r*’ is known.

## BitterMatch Algorithm

### Feature extraction from similarities

The ligand similarity matrices determine the neighbors (or high similarity ligands) of a given ligand. We developed neighbor-informed features that summarize the abundance of positive and negative associations in the neighborhood of a given ligand to a receptor of interest. From each ligand similarity matrix, we annotate each ligand (*l*)-receptor (*r*) pair with four features: two summarize the similarities of the ligand *l* to ligands with *positive associations* to the receptor *r* and two summarize the similarities of *l* to ligands with *negative associations*. We measure the similarity to the nearest ligand that activates *r*:

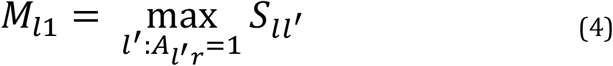

and the sum of similarity values to all ligands that activate receptor *r*

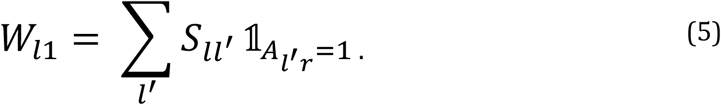

Similarly, we measure the similarity to the closest ligand that does not activate receptor r:

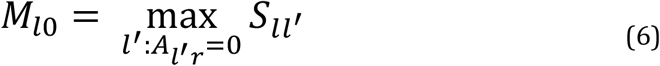

and the sum of similarity values to all ligands that do not activate receptor *r*:

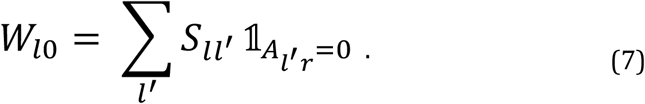

Likewise, we extract neighbor-informed features from each receptor similarity matrix, reversing the roles of ligand and receptor. We measure the similarity to the closest receptor that is activated by the ligand *l*:

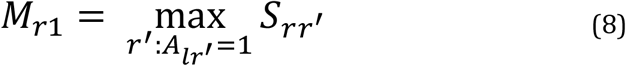

and the sum of similarities to all receptors that are activated by the ligand *l*:

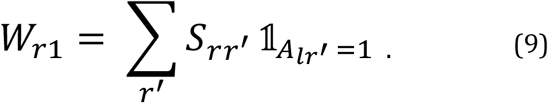

Similarly, we measure the similarity to the closest receptor that is not activated by ligand *l*

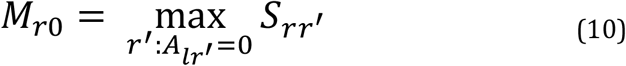

and the sum of similarities to receptors that are not activated by the ligand *l*:

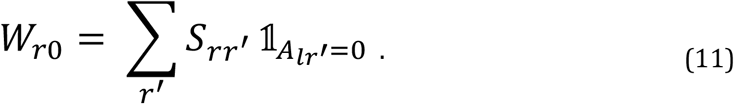

### Feature matrices

We construct a feature matrix *X* ∈ ℝ^*RL×m*^ in which each row corresponds to a ligand-receptor pair, and each column corresponds to a feature. We denote by *m* the number of features. All models include *C_l_=235* chemical descriptors of the ligand (Methods: “Ligand features”) and *C_r_=250* chemical descriptors of the receptor (Methods: “Receptor features”). The models differ by the number of similarity-based features included. Models 2, 3, 4 and the “new ligands” model include four features for each type of similarity as described in Methods: “Ligand similarities”, “Receptor similarities and identities”. Correspondingly, we construct a label vector *Y* ∈ ℝ^*RL*^ by vectorization of the association matrix *A.*

We construct train and test feature matrices *X*^Tr^,*X*^Te^ and label vectors *Y*^Tr^,*Y*^Te^ by randomly partitioning our data into a training set and a test set. The sampling units are matched to the scenario. In the *“filling the gaps”* scenario, we sample individual cells (i.e. ligand-receptor pairs), in a balanced manner. That is, we sample ρ=0.8 of the positive associations and ρ=0.8 of the negative associations within each receptor into the training set, and the rest form the test-set. In the *“new ligands”* scenario, we sample ρ=0.8 of the ligands for the training set, meaning that the training set is composed of their known positive and negative associations with all the receptors.

To avoid data leakage, before constructing the collaborative similarity matrices, we treat the values corresponding to test pairs as missing values in the association matrix *A.*

### Classification

We model the problem of predicting the TAS2Rs that associate with a ligand as a binary classification task where each ligand-receptor pair is given a prediction.

We train a gradient boosting classifier with decision tree learners using the XGBoost classifier on the train feature matrix *X*^Tr^ and the corresponding train label vector *Y*^Tr^ to predict the outcome for each ligand-receptor pair. The hyper-parameters used for the XGBoost classifier were not chosen using the data, and are detailed in the Table S2.

## Evaluation

### Baseline models

The sub-models of BitterMatch were compared with a prior model that represents a naive guess. The prior model gives a fixed prediction *ϕ_r_* for all ligands associating with receptor *r*. For estimation of *ϕ_r_* we use the proportion of ligands that activate the receptor *r* in the training set. Note that *ϕ_r_* does not depend on the ligand, but only on the receptor.

In the *“New ligands”* scenario, BitterMatch is also compared with a nearest-neighbor model according to the linear similarity. Denote by

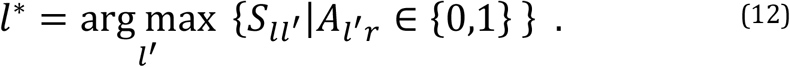

the nearest ligand to *l*, whose association with *r* is known. Then the nearest-neighbor model uses the association of the nearest ligand as *A_l*r_* its prediction for the ligand *l*. Equivalent results were achieved when predictions are given according to MOLPRINT2D similarity (not shown).

### Recall-Precision

In order to evaluate performance of BitterMatch models (described in Methods: “Filling the gaps scenario” and Methods: “New ligands scenario”), we performed 100 repetitions of the experiment. In each repetition, train and test feature matrices *X*^Tr^,*X*^Te^ and label vectors *y*^Tr^,*y*^Tr^ were randomly sampled (as described in Methods: “Feature matrices”).

For each model we calculated an average precision-recall curve over 100 repetitions of the experiments. Denote by tp the number of true positive predictions, by *fp* the number of false positive predictions, and by *fn* the number of false predictions. The precision levels were calculated as 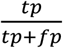 and recall levels were calculated as 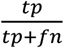. The average precision scores were computed as 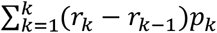, where *p_k_* and *r_k_* are the precision and recall at the *k*-th threshold. The number of considered thresholds *K* was set to the number of unique prediction scores.

Since in each repetition we observe the precision at a different set of recall values, we use linear interpolation to a fixed set of locations. We set *u*_1_,..., *u_k_* as k = 200 fixed levels of recall, evenly spread between 0 and 1. For each repetition *i*, we treat the precision values 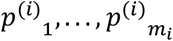 as a function of the recall values 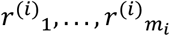 and used these values to perform linear extrapolation of the function, evaluated at *u*_1_,...,*u_k_*. Having 100 sets of precision values, each evaluated at the same recall levels *u*_1_,..., *u_n_*, we calculated asymptotic 95% confidence intervals for the mean precision at each recall level 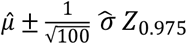 where 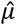 is the mean precision at the given recall level, 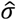 is the sample standard deviation of the precision values at the given recall level over 100 repetitions, and *Z*_0.975_ the critical value of the standard normal distribution. To obtain prediction bands we set the lower and upper bounds of the prediction bands at *u_j_*, as the 5-th and 95-th percentiles of *p*^(*i*)^_*j*_. Percentiles were computed using the “percentile” function, and the interpolation was performed using the “interp” function, both from the numpy 1.16.4 Python library.

### Feature importance

Average gain across all XGBoost splits was computed as defined in^37^ using the “get_booster” function (with importance_type=‘gain’) from the xgboost 1.4.2 Python library.

## Validation set

### Curation of molecules for validation set

The validation set was created with new compounds that were neither used for training nor testing in the experiment. In total, 12 compounds were collected: 3 compounds that were not tested *in-vitro* to the best of our knowledge, 4 compounds taken from the work of Lang et al.^41^ and 5 compounds from BitterDB that were not included in the original association matrix (only one or two known associations are documented for them in BitterDB). In total, 252 pairs of ligand-receptor were predicted for this set. The evaluation was performed at a threshold of 0.52 was chosen on the test set.

## *In-vitro* testing of BitterMatch predictions

### Chemicals and Materials

The compounds for functional analyses were purchased from Alfa Aesar (Ward Hill, USA) (2-acetylbenzofuran, fisetin), Sigma-Aldrich (Steinheim, Germany) (apigenin, butein, 3,2’-dihydroxycalchone, quercetin and sinapic acid) and AdipoGen (Liestal, Switzerland) (theacrine), dissolved in dimethyl sulfoxide (DMSO) according to their solubility and stored at −20 °C until use.

### Functional receptor screening

HEK 293T-Gα16gust44 cells^41^ were seeded into 96-well-plates and cultivated in DMEM plus supplements (10% FCS, 1% penicillin/streptomycin, 1% glutamine) at 37 °C, 5% CO2, saturated air humidity. Next, cells were transiently transfected with cDNA constructs coding for the 25 human TAS2Rs using lipofectamine 2000 (Thermo Fisher Scientific GmbH, Darmstadt, Germany) as before^41,62^. An empty vector (mock) was transfected as a negative control. After 24 h, cells were loaded with Fluo4-AM (Thermo Fisher Scientific GmbH, Darmstadt, Germany) in the presence of 2.5 mM probenecid, washed two times for 30 min. with C1 buffer (130 mM NaCl, 5 mM KCl, 2 mM CaCl2, 10 mM glucose, 10 mM HEPES; pH 7.4) and placed in a fluorometric imaging plate reader (FLIPRTetra, Molecular Devices). For screening, stock solutions of the compounds were diluted in C1 buffer. The final DMSO concentration did not exceed 1%. The final maximal concentrations of the compounds added to the receptor-transfected cells were between 1 and 1000 μM depending on their solubilities and the occurrence of receptor-independent cellular artefacts (Supplementary Table S1). Additional to the maximal compound concentration, a tenfold dilution of the maximal concentration were tested to judge the apparent potencies. As positive controls we included the known agonists aristolochic acid (1 and 10 μM, TAS2R14)^63^ and strychnine (30 and 300 μM, TAS2R10)^64^ and (3 and 30 μM, TAS2R46)^65^. The changes in fluorescence upon addition of compounds were monitored. Vitality of cells was confirmed by the addition of somatostatin 14 (100 nM, Bachem, Bubendorf, Switzerland) as a second application. Screening experiments were performed in duplicate wells. A statistically significant difference between the fluorescence changes of receptor-transfected cells compared to mock-transfected cells was judged as potential hit and the corresponding compound-receptor pairs were selected for further experiments.

### Determination of threshold values of TAS2Rs activation

The evaluation of receptor-compound pairs judged positive after the initial screening was based on three additional independent experiments. For the calculation of ΔF/F, the fluorescence change of control (mock) cells was subtracted from that of the corresponding receptor-expressing cells. The resulting signal was normalized to background fluorescence.

### Functional analogs analysis

Prediction of functional orthologs was performed on the full binary association matrix containing both the original data and the predictions (threshold = 0.65) in cases where the associations were unknown. We calculated the Jaccard similarity between human and mice receptors, taking into account intersections in both positive and negative associations.

### Prediction of DrugBank dataset

We used BitterMatch to predict the associations of intensely bitter drugs from DrugBank (version 5.1.5)^42^. 2406 intensely bitter predicted drugs were taken from Margulis, E. et al.^16^ and inputted into BitterMatch. Analysis was performed at a threshold of 0.52.

## Implementation

All the code for this work was implemented in Python 3.6. For the XGBoost implementation we used the xgboost 1.4.2 Python library. Figures 2, 4, 6, S1, S2, S3, S4 were generated with matplotlib 3.3.4. Figures 3,5 were generated with matplotlib 3.2.2. Figure 4B was implemented using seaborn 0.9.0. Figure 1 and S1 were Created with BioRender.com.

## Code availability

The code is available at: https://github.com/YuliSl/BitterMatch

## Acknowledgements

We thank Lior Wolf, Ayana Dagan-Wiener and Fabrizio Fierro for insightful discussions, and Eva Boden for excellent technical assistance. MYN is supported in part by ISF #1129/19 funding. EM is the recipient of CIDR, Smith, Zehavi and Nano fellowships. YS is supported by the Israeli Council For Higher Education Data Science fellowship.

## Authors contributions

MYN conceived the study, EM, YS, YB and MYN designed the study and wrote the manuscript. EM designed the data set and features, YS developed the learning algorithm. EM and YS analyzed the results. TL and MB performed in-vitro experiments. All authors read and approved the manuscript.

## Supplementary data

**Figure S1.**
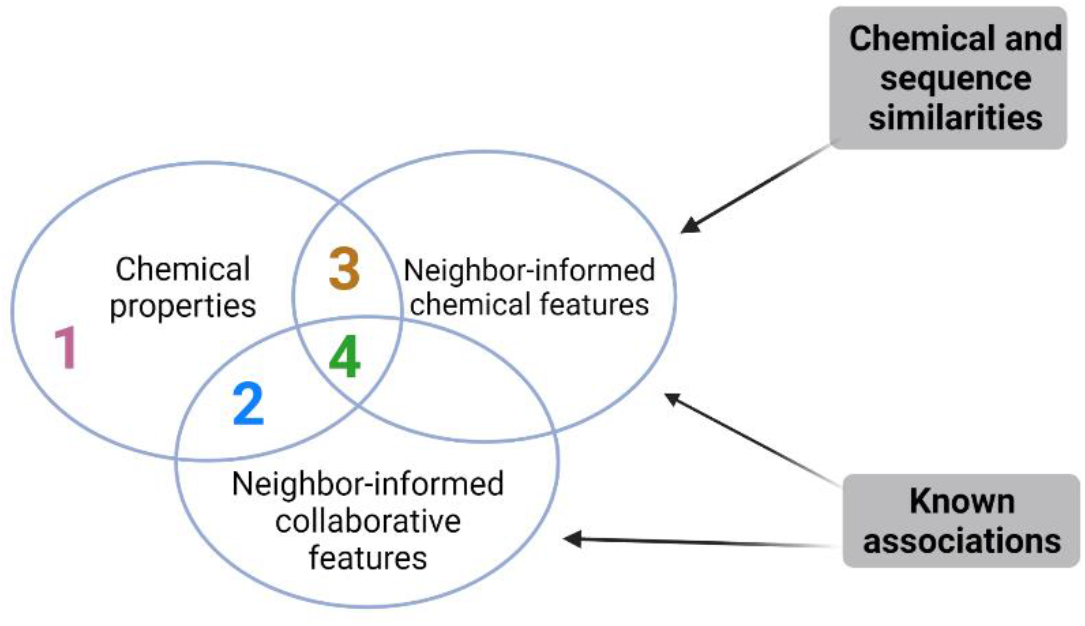
BitterMatch sub-models. A Venn diagram describing the four examined BitterMatch sub-models. Model (1) includes only chemical properties, model (2) includes chemical properties and neighbor-informed collaborative features, model (3) includes chemical properties and neighbor-informed chemical features, model (4) is the augmented one and it includes chemical properties, neighbor-informed collaborative features and neighbor-informed chemical features. Neighbor-informed collaborative features are computed directly from the known associations that were also used to calculate collaborative similarities. However, neighbor-informed chemical features are computed from the known associations and chemical and sequence similarities.

**Figure S2.**
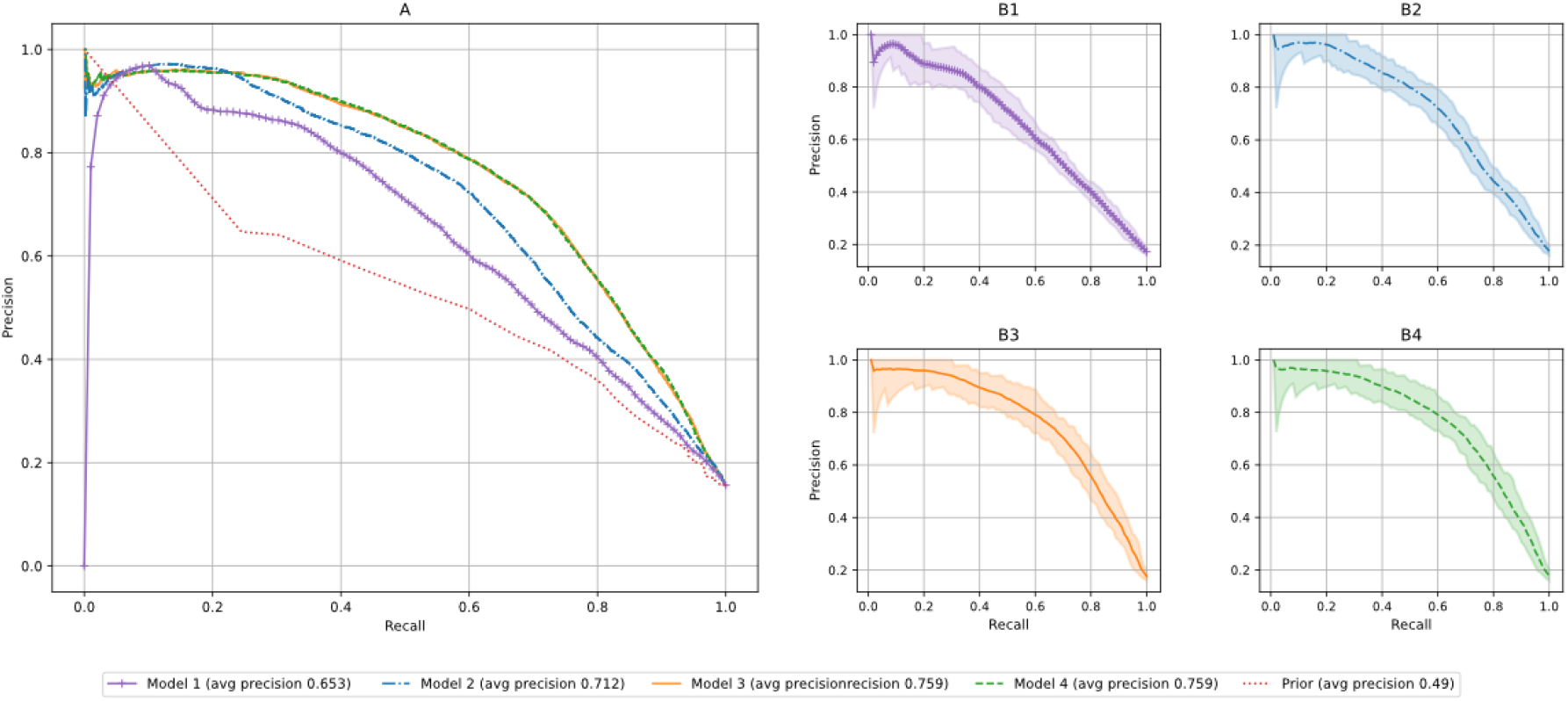
Prediction intervals for filling the gaps scenario. Average precision-recall curves over 100 repetitions are shown in (A) for the four BitterMatch models and for the prior model (identical to Figure 2A). Figures B1, B2, B3, B4 show Bootstrap 95% prediction intervals for each model presented in (A).

**Figure S3.**
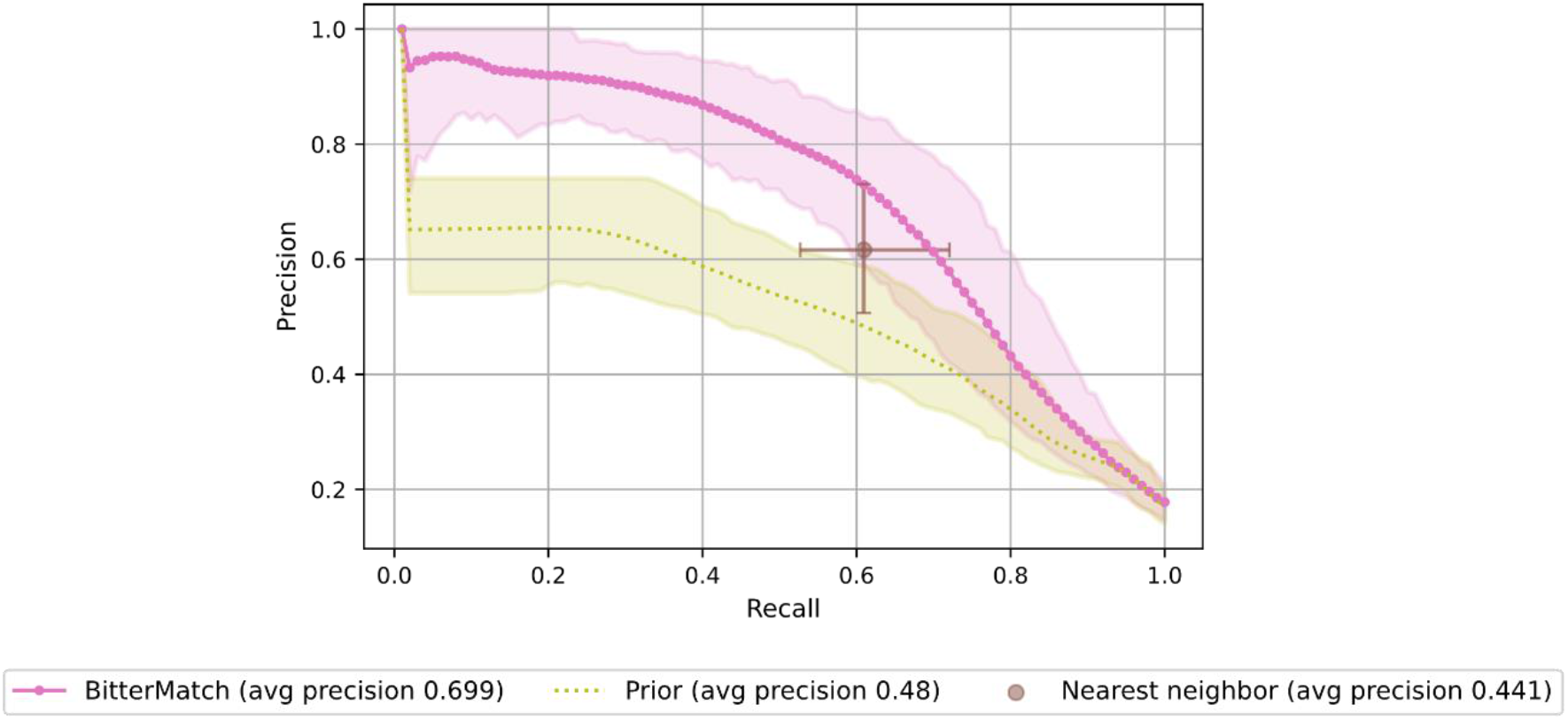
Prediction intervals for new ligands scenario. Precision-recall curves for the adapted BitterMatch model, a prior model and a nearest neighbor model (idenctivcal to Figure 4A). 95% Bootstrap prediction intervals are shown for each model.

**Figure S4.**
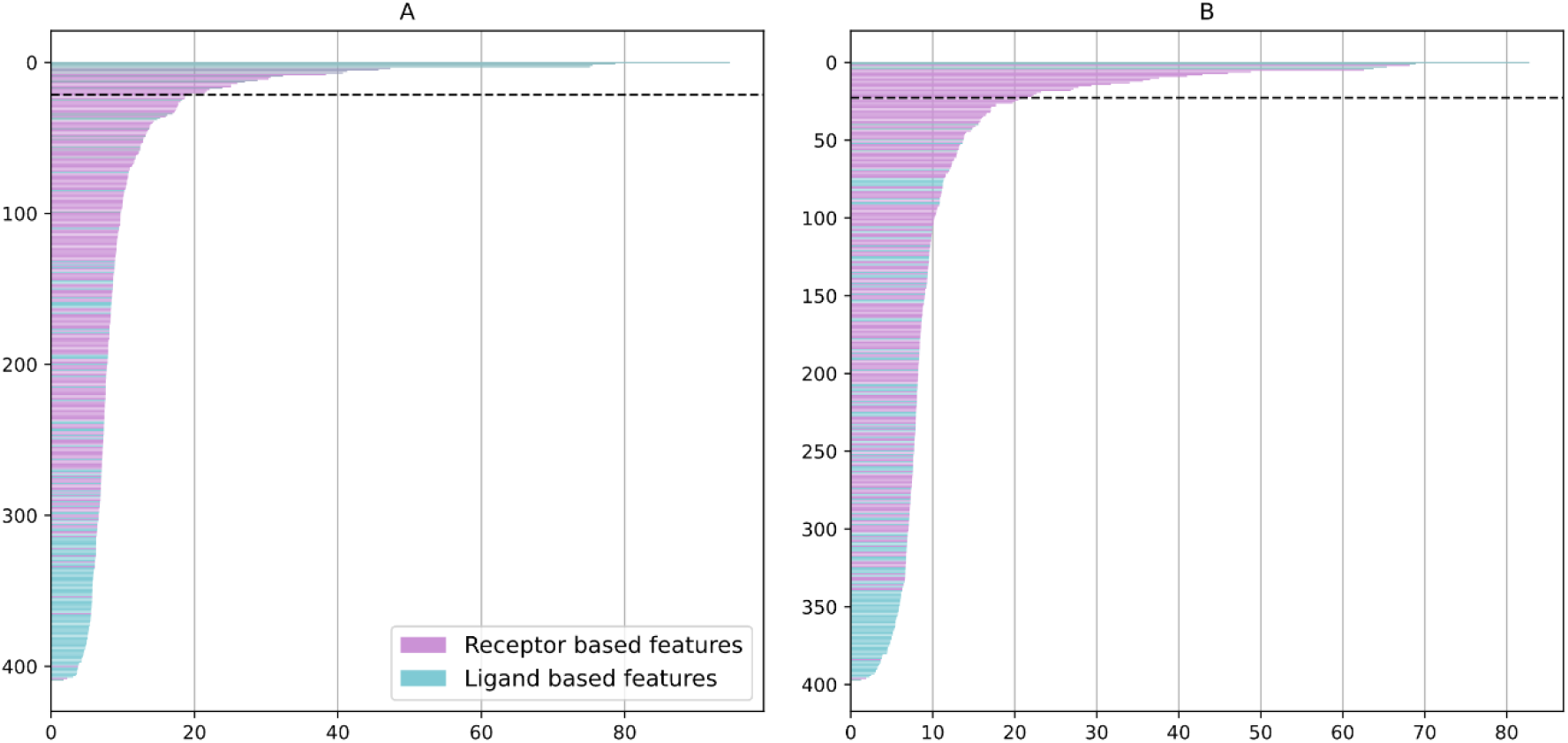
Feature importance for new ligands. Features related to ligand similarities are shown in blue, receptor properties are shown in purple. (A) – Feature importance for all the features in model 3. The black horizontal dashed line corresponds to feature importance of 21.5, features above this threshold are shown in detail in Fig. 6 (B) – Feature importance for all the features in the “new ligands” model. The black horizontal dashed line corresponds to feature importance of 22.7.

**Figure S5.**
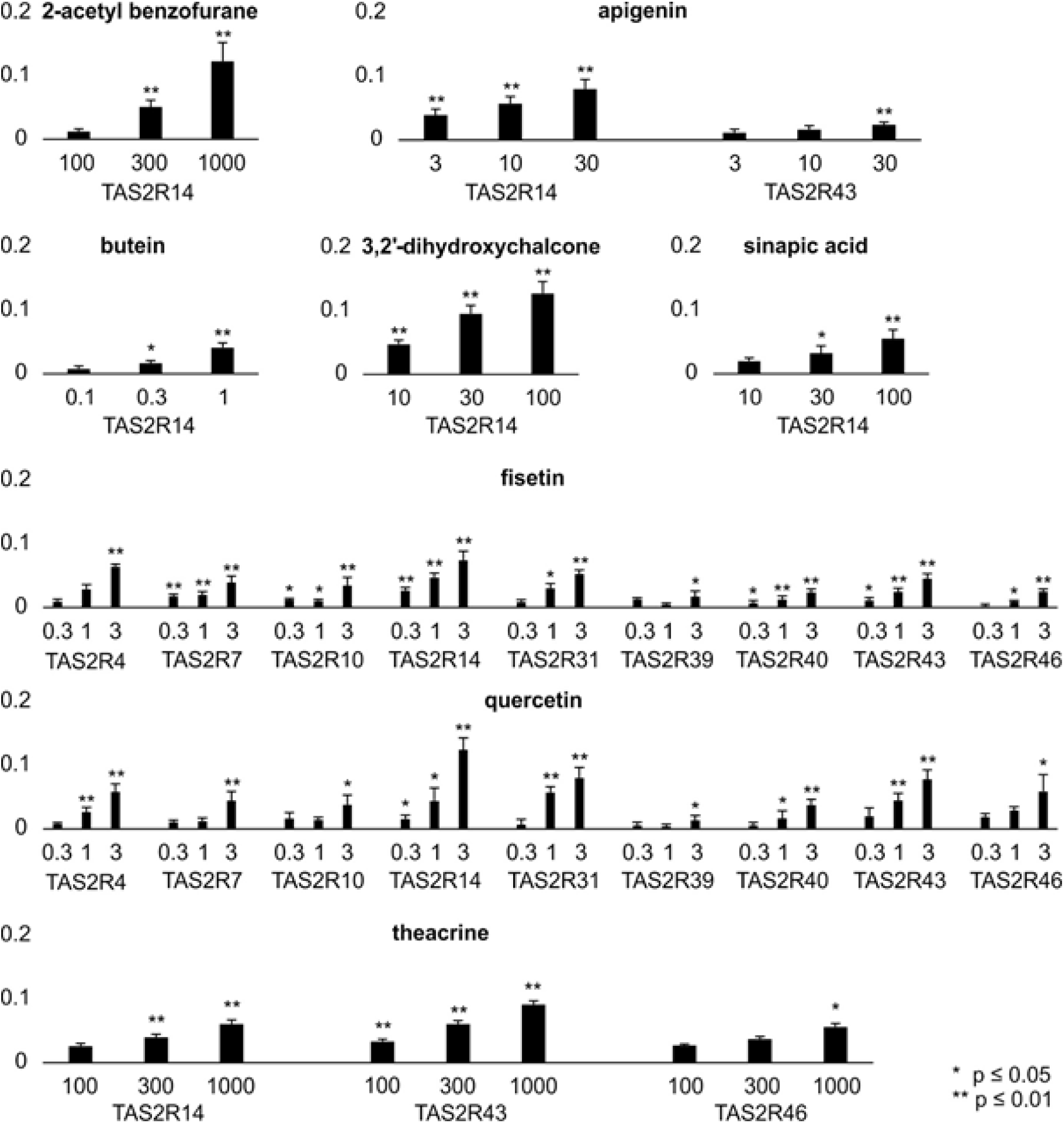
Confirmation of TAS2R screening results by functional experiments. The cDNA of the 9 TAS2Rs showing responses to the predicted bitter substances were expressed in HEK 293T-Ga16gust44 cells and challenged with 3 concentration of the corresponding newly identified agonists. The relative changes of fluorescence (ΔF/F) are provided on the y-axes of the bar graphs for each substance. Below the bars the concentrations used for the confirmatory experiments in μM are depicted together with the corresponding receptor symbols. The substance names are printed in bold. Statistically significant activations (Student’s t-test) are indicated by asterisks (see bottom right of figure for significance levels).

**Table S1.**
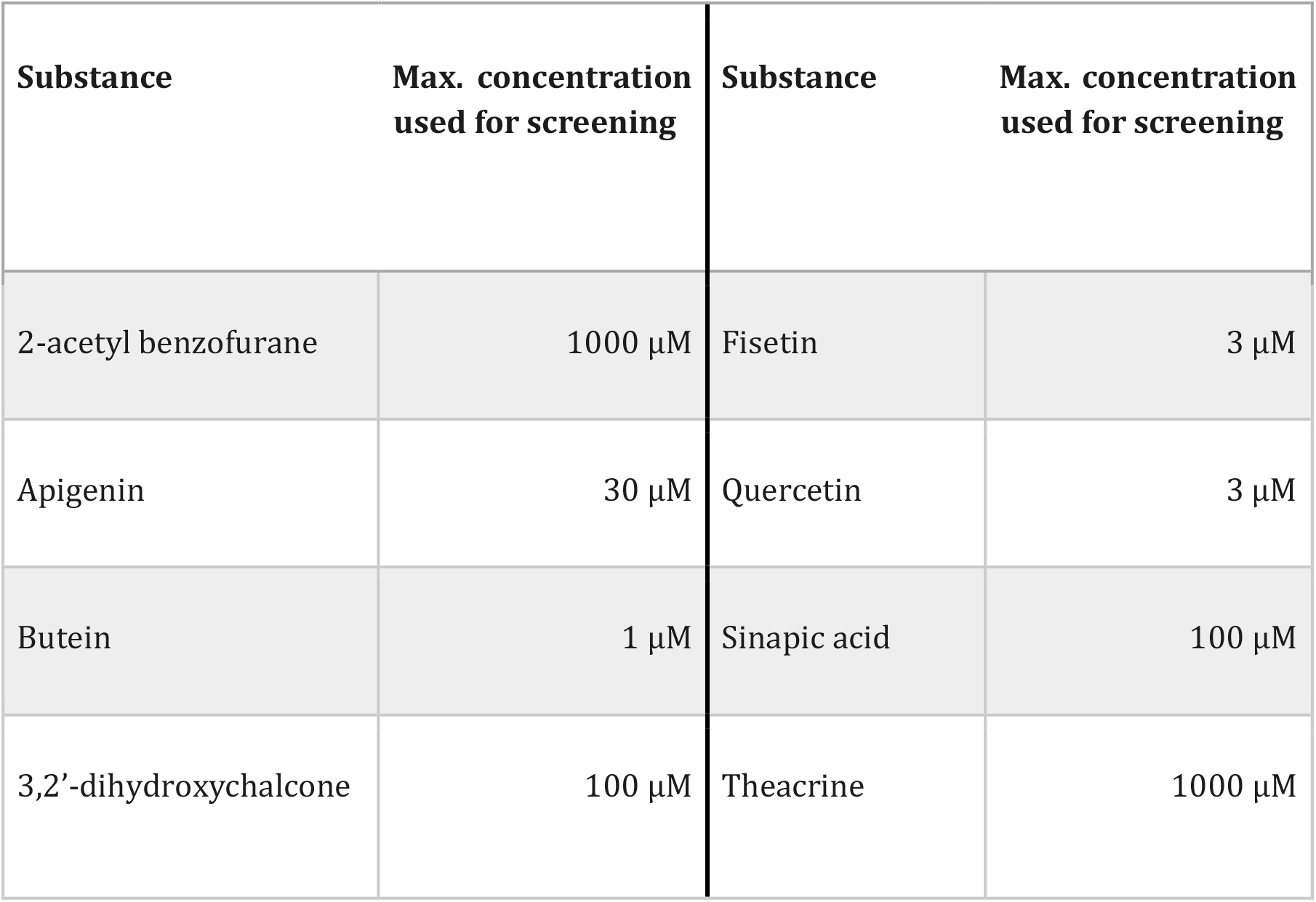
Substances used for experimental testing of prospective predictions and their maximal employed concentrations.

**Table S2.** XGBoost hyper-parameters. To avoid overfitting 1000 trees were used accordingly the following hyper-parameters were adjusted. The rest of the parameters were set to their default values.

| Parameter | Value |
| --- | --- |
| Number of estimators | 1000 |
| Learning rate | 0.001 |
| Maximal depth of a tree | 4 |
| Column sample by tree | 0.3 |
| $\gamma$ - minimum loss reduction for partition on a leaf of a node | 2 |
| Minimum sum of instance weight needed in a child | 0.45 |
| Subsample - ratio of the training instances used prior to growing trees | 0.7 |

**Table S3.**
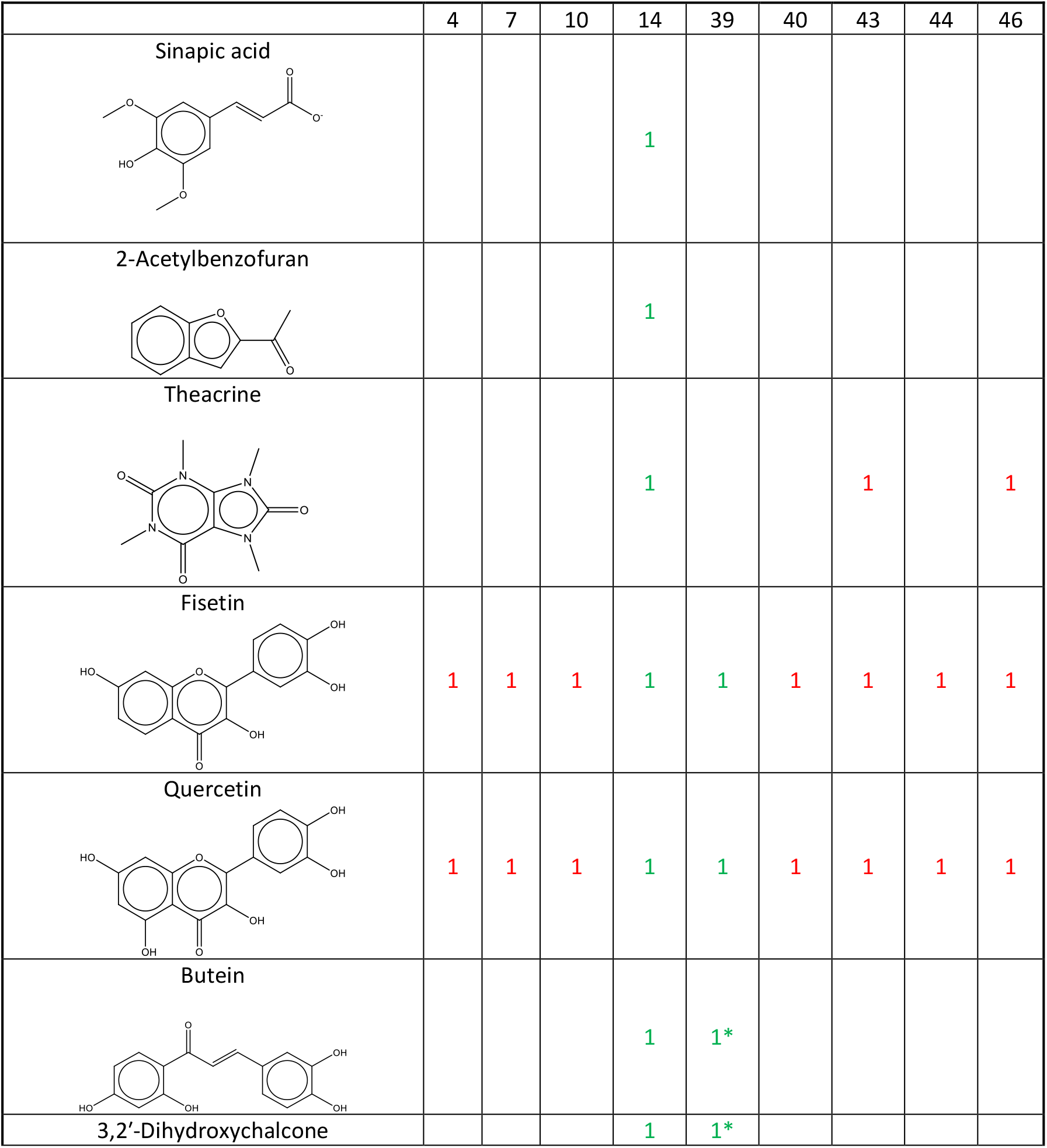
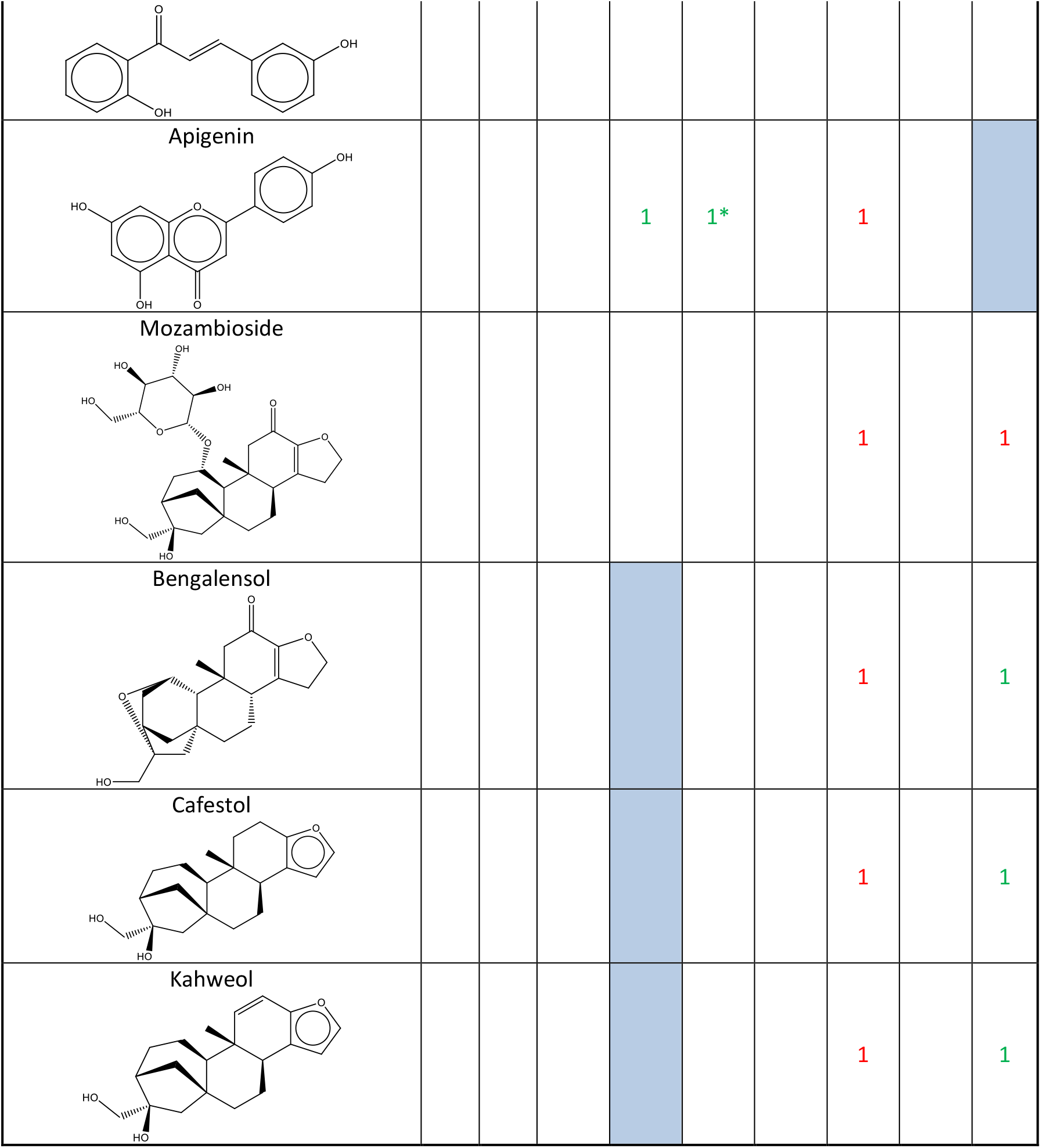
Prospective prediction results per ligand. 1 –represents an activation of the receptor that was confirmed experimentally and blank space means no activation was detected. * – positive activation that was detected in another publication but not in our in-vitro experiment. 1–TP, 1-FN, blue-FP and White – TN

|  | 4 | 7 | 10 | 14 | 39 | 40 | 43 | 44 | 46 |
| --- | --- | --- | --- | --- | --- | --- | --- | --- | --- |
| **Sinapic acid** |  |  |  | 1 |  |  |  |  |  |
| **2-Acetylbenzofuran** |  |  |  | 1 |  |  |  |  |  |
| **Theacrine** |  |  |  | 1 |  |  | 1 |  | 1 |
| **Fisetin** | 1 | 1 | 1 | 1 | 1 | 1 | 1 | 1 | 1 |
| **Quercetin** | 1 | 1 | 1 | 1 | 1 | 1 | 1 | 1 | 1 |
| **Butein** |  |  |  | 1 | 1* |  |  |  |  |
| **3,2'-Dihydroxychalcone** |  |  |  | 1 | 1* |  |  |  |  |
| Apigenin |  |  |  | 1 | 1* |  | 1 |  |  |
| Mozambioside |  |  |  |  |  |  | 1 |  | 1 |
| Bengalensol |  |  |  |  |  |  | 1 |  | 1 |
| Cafestol |  |  |  |  |  |  | 1 |  | 1 |
| Kahweol |  |  |  |  |  |  | 1 |  | 1 |

